# Approximation of the age distribution of cancer incidence using a mutational model

**DOI:** 10.1101/2021.07.06.451349

**Authors:** Alexandr N. Tetearing

## Abstract

The approximation of the age distributions of cancers was carried out using a complex mutational model presented in our work [1]. Datasets from the American National Cancer Institute (SEER program) were used.

We approximated the datasets for the age distributions of lung, stomach, colon and breast cancer in women; cancer of the lung, stomach, colon and prostate in men.

The average number of mutations (required for cancer formation) averaged over the four types of cancer is 5 mutations per cell in women and in men. The average (over the four types of cancer) mutation rate is estimated as 1.0 · 10 ^−1^ mutations per year per cell for women and 3.8 · 10 ^−1^ mutations per year per cell for men.

This article is a continuation of work [1].

## 1. Mutational model of age distribution of cancers

In this article, we use the Dz(t) function obtained in [1] as a fitting function to approximate the datasets on the age distribution of cancer incidence:

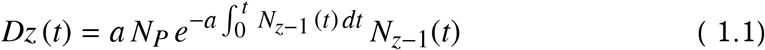

Here *z* is the number of cell mutations required for cancer to occur.

*N*_*P*_ is the number of people in the considered group (the group with the given age distribution of cancers).

The function *N*_*z*−1_ (*t*) describes the number of cells (in the cell population of anatomical tissue) with the number of mutations (*z* − 1) or more:

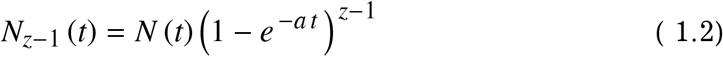

The *a* parameter is the average number of mutations (key events) that occur in a cell per unit of time.

The *N*(*t*) function is the growth function of the cell population (anatomical tissue). The function describes the number of cells in the cell population of the organ (tissue). The growth of the cell population occurs in accordance with the equations:

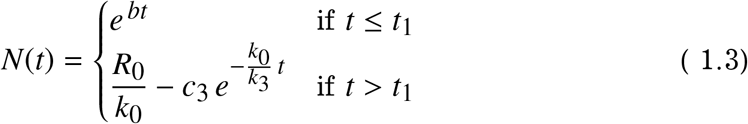

Until time *t*_1_ the population (number of cells) grows exponentially (the equation of unlimited growth), after the time *t*_1_ the population is faced with a limitation in the food (energy) resource (the equation of limited growth).

Thus, the time *t*_1_ is the time when the cell population has the maximum growth rate.

The constant *b* from the upper equation of system (1.3) is determined by the expression:

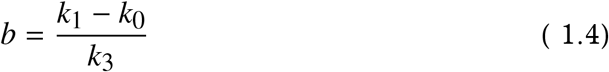

The coefficients *k*_*i*_ are constant coefficients that characterize the interaction of cells with the environment.

The constant *c*_3_ (this is the constant of integration) is found from the equation:

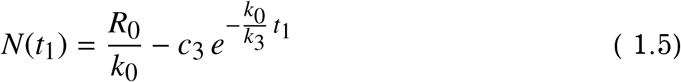

The *R*_0_ constant determines the amount of resource available to the cell population per unit of time (at the stage of limited growth). The *R*_0_*/k*_0_ ratio determines the number of cells in the population at the final stage of population growth.

The time *t*_1_ (the time when the cell population passes from the stage of unlimited growth to the stage with limited growth) is determined by the equality:

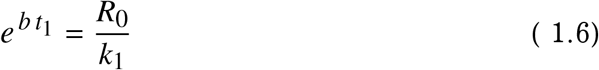

All these equations and coefficients are described in detail in book [2].

The *N*(*t*) function is shown in Fig. 1.1.

**Figure 1.1.**
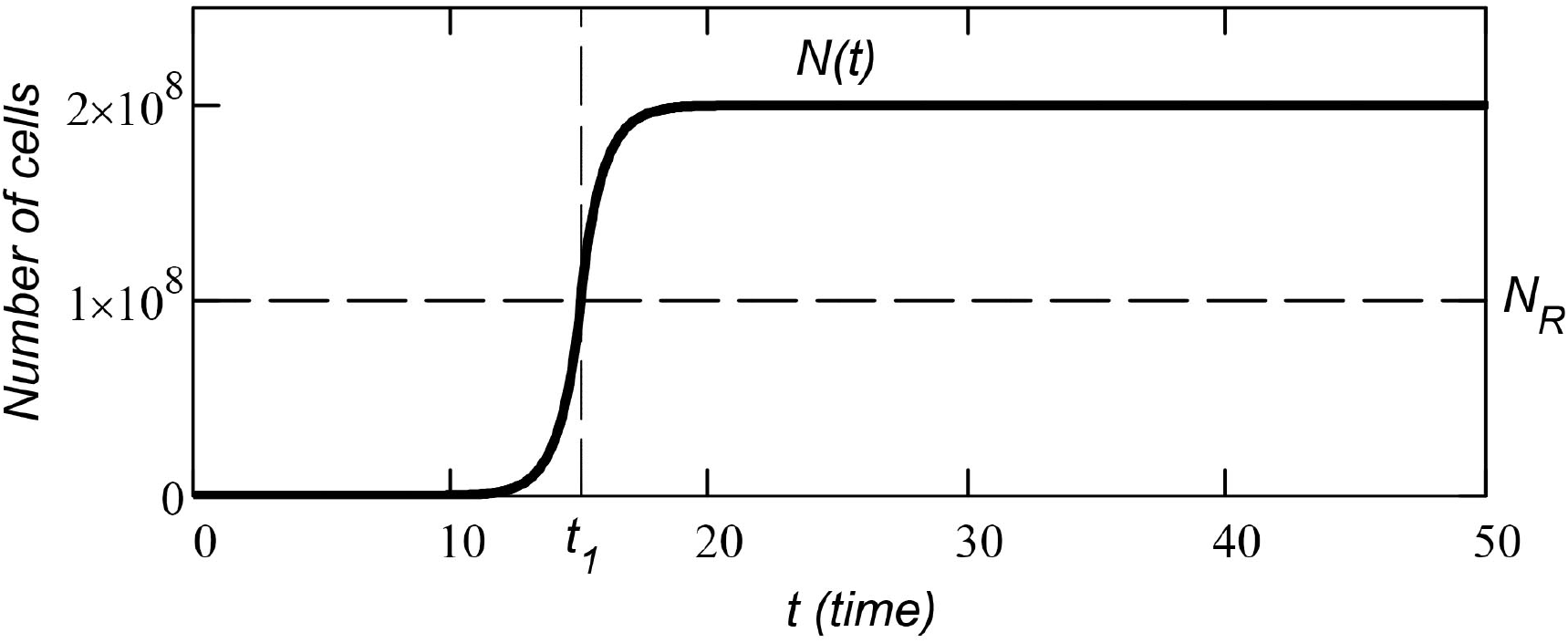
Growth function of the cell population of anatomical tissue.

For simplicity, we assume that at time *t* = 0 the cell population consists of one cell: *N*(0) = 1. When plotting the graph in Fig. 1.1 we used the following parameters: *k*0 = 0.5, *k*1 = 1, *R*0*/k*0 = 2.0 · 10 ^8^, *t*1 = 15.

## 2. Datasets

We used data from the SEER (Surveillance Epidemiology and Outcomes Program) of American National Cancer as the data source [3].

These data are more detailed in comparison with the generalized data of the Belorussian register of cancers, which we used earlier in [1].

SEER data cover the age range of patients from 0 to 110-115 years. The age of initial cancer registration is recorded for all patients. Thus, we can obtain the age distribution of the incidence by single year of age.

For example, in the datasets provided by SEER, there are cases where lung cancer in women was registered at the age of 113, breast cancer at 118; in men, there is a case when lung cancer was diagnosed in a patient at the age of 111 years.

However, the age distribution of the U.S. population (census data) is given by single year of age only for the age range from 0 to 99. Data for ages over 99 grouped into one number and therefore cannot be used as data to find the fitting function (see next section).

We used data on all registered patients (separately for female and male cancers) registered from 2000 to 2009. All patients enrolled in the SEER program (patients aged 0 to 99) are considered without racial separation. The total number of cases considered is:

### Women

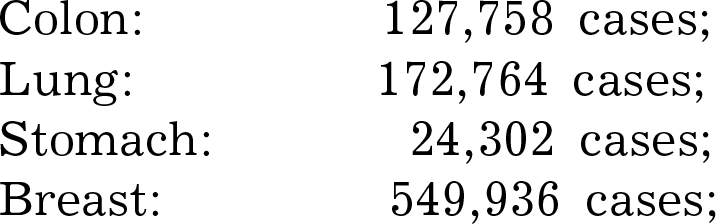

|  |  |
| --- | --- |
| Colon: | 127,758 cases; |
| Lung: | 172,764 cases; |
| Stomach: | 24,302 cases; |
| Breast: | 549,936 cases; |

### Men

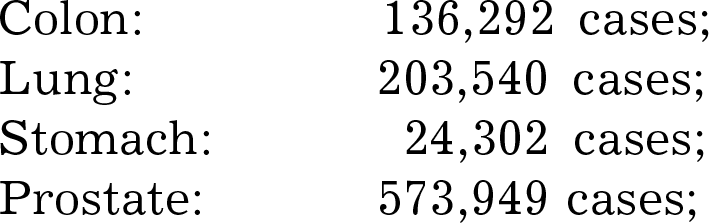

Men:
|  |  |
| --- | --- |
| Colon: | 136,292 cases; |
| Lung: | 203,540 cases; |
| Stomach: | 24,302 cases; |
| Prostate: | 573,949 cases; |

## 3. Data preparation

In order to plot the age distribution of incidence, it is necessary to recalculate the baseline data on reported cases of cancer, taking into account the distribution of the population by age groups. Fig. 3.1 shows an example of such a recalculation of data (for the year 2000).

**Figure 3.1.**
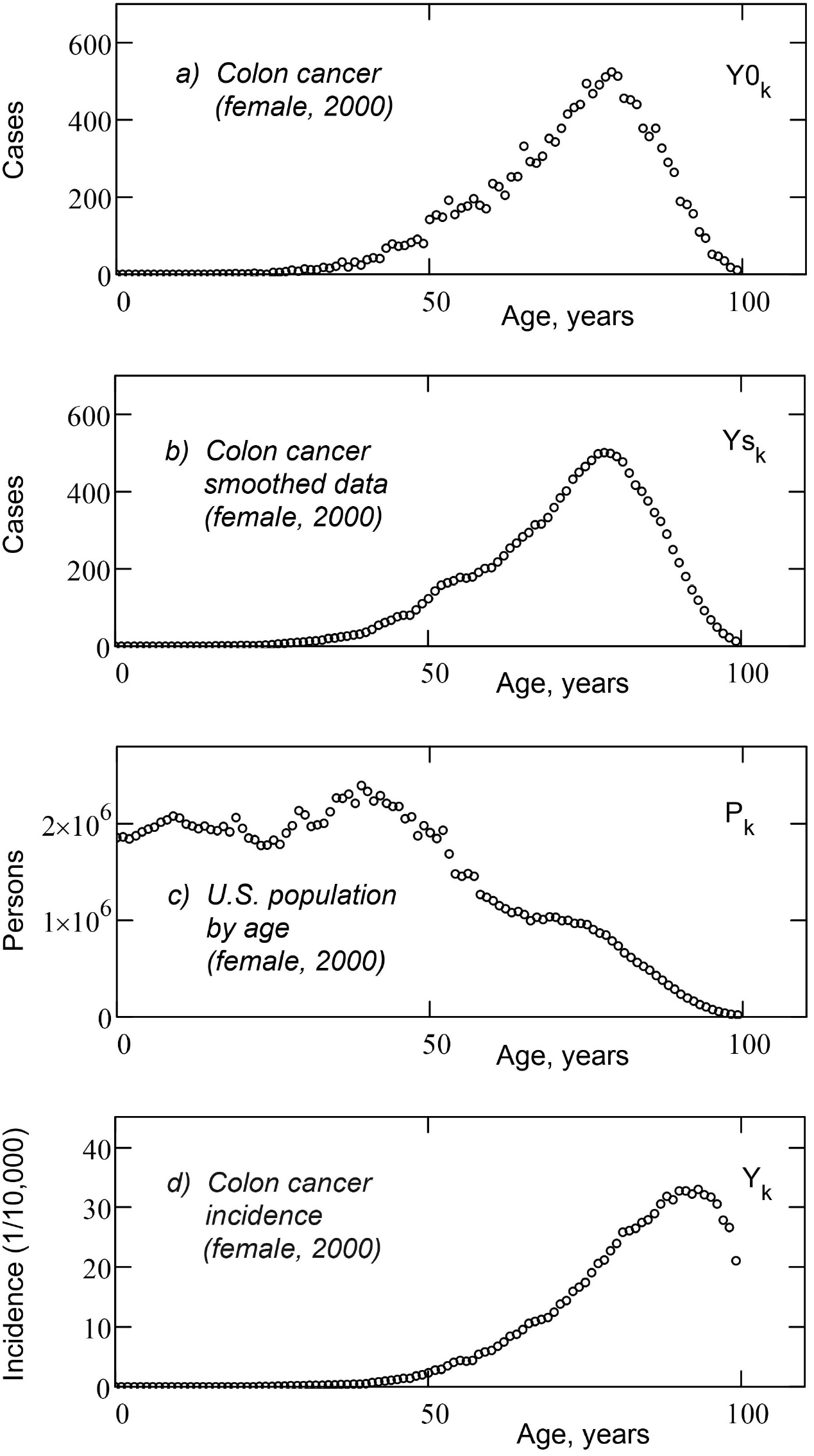
Preparing data for approximation: a) initial data (cases registered in 2000); b) smoothed data; c) distribution of women by age in 2000); d) age distribution of colon cancers in women per 10,000 people (for 2000).

The top graph (Fig. 3.1-a) shows the initial data (registered cases of colon cancer in 2000, distributed by patient age). The graph is a discrete dataset *Y*0_*k*_, where *k* = 0, 1, 2 … 98, 99 (age of patients).

Fig. 3.1-b shows the smoothed data 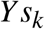 (median smoothing with 5-year bandwidth).

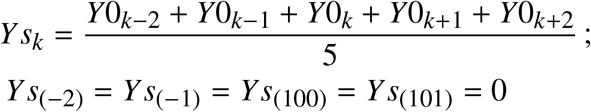

Smoothing (without reducing the total number of cases) reduces the scatter of the data when the number of cases is small.

Recall that these are cases registered only in 2000 (and we cannot add here for averaging the cases registered, for example, in 2001, since in 2001 the population has a different age distribution).

Figure 3.1-c shows the age distribution of female population for U.S. in 2000 (*P*_*k*_ data set).

The graph in Fig. 3.1-d (*Y*_*k*_ incidence data) is obtained as the quotient of dividing graph 3.1-b by graph 3.1-c, multiplied by 10,000 and divided by 0.28 (as SEER data covers only 28% of the U.S. population [4]).

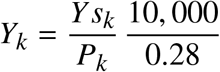

As a result, we get the age distribution of cancers (colon cancer in women) per 10,000 people in 2000, which is shown in Fig. 3.1-d.

Comparing the data in Figs 3.1-b and 3.1-d, we see that the graphs are very different. The shape of the distribution has changed, and the maximum of the graph has shifted 15 years to the right.

After doing the described procedure 10 times for each calendar year (from 2000 to 2009), we sum up the ten obtained annual distributions and get the age distribution of cancers per 100,000 people, averaged over 10 years (for the final graph, see the section “Results”, the top graph in Fig 5.1).

**Figure 5.1.**
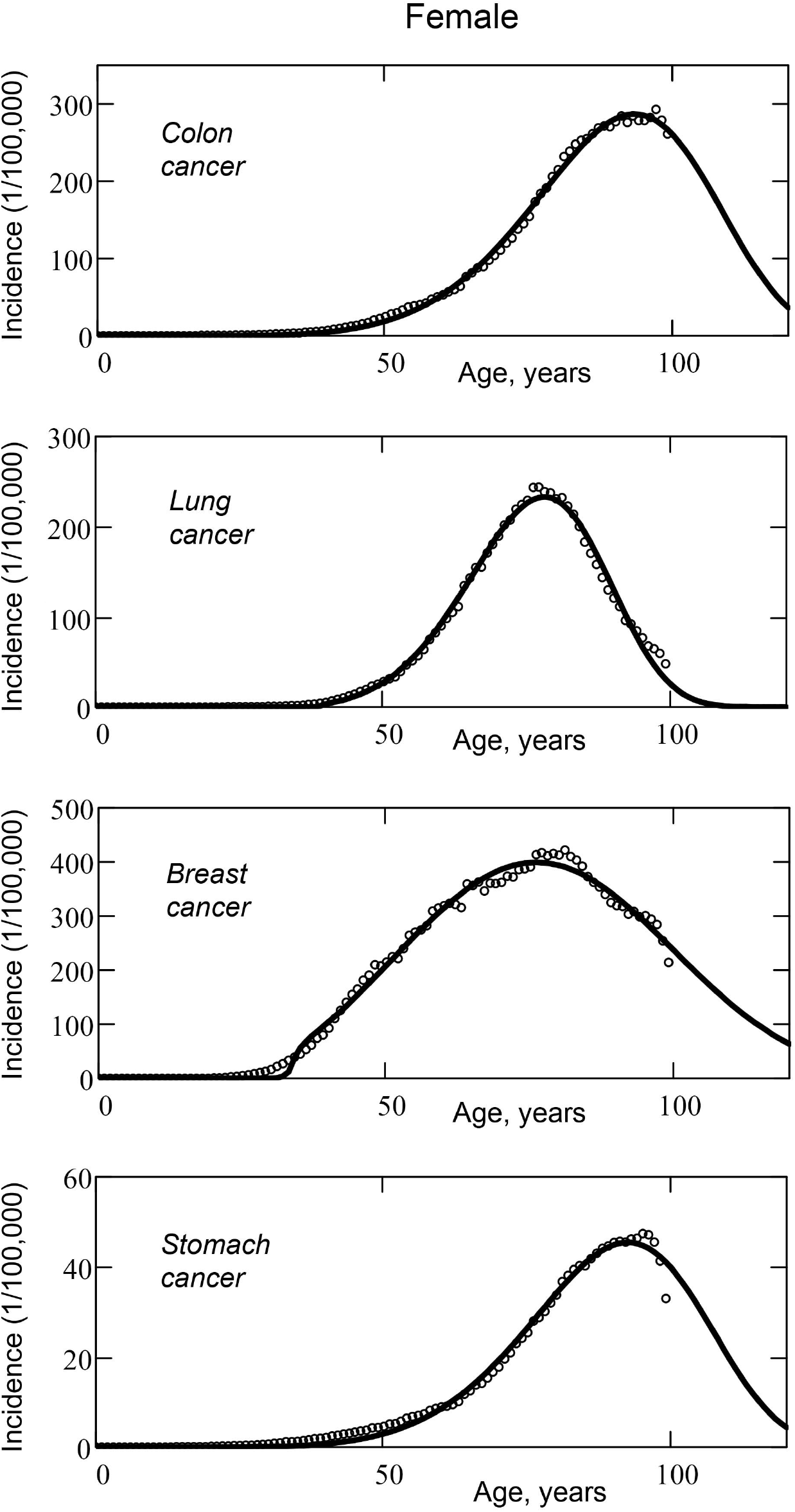
Age distribution of cancer incidence in women. The light points are real datasets; solid bold curves are approximation functions (mutational model of cancer).

The data obtained in this way are ready for approximation by the model function of age distribution.

## 4. Approximation method

As an approximating function, we used function (1.1), which describes the mutational model of cancer, presented in [1].

As in [1], the optimization of the error function *Er* is carried out programmatically, by the conjugate gradient method for each fixed value of *z*, since the number of mutations *z* (the number of key events required for the onset of cancer in a cell) is a positive integer.

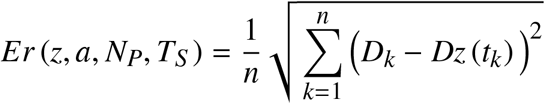

Here *t*_*k*_ is the age from the real dataset; *D*_*k*_ is the value of the age distribution from the real dataset for the age of *t*_*k*_; *Dz* (*t*_*k*_) is the theoretical value of the age distribution for the age *t*_*k*_.

The function is optimized by four parameters: *z, a, N*_*P*_ and *T*_*S*_. We select a set of parameters for which the error function *Er* (*z, a, N*_*P*_, *T*_*S*_) is minimal (least squares method).

We also optimize the delay parameter *T*_*S*_ added to formula (1.1) – we assume that cancer is diagnosed not at the moment of onset, but several years later (after the time *T*_*S*_). That is, the time *T*_*S*_ must not be negative or equal to zero. Therefore, we substitute the difference (*t* − *T*_*S*_) into the equation for *Dz*(*t*) function instead of time *t*.

When approximating, we used the following constant parameters: *k*_0_ = 0.5; *k*_1_ = 1; *t*_1_ = 15. The coefficient *k*_3_ is calculated from the equation:

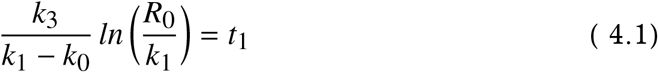

The *R*_0_ parameter is chosen (for each organ) so as to provide the required number of cells at the final stage of organ growth, which is determined by the anatomical size of the organ. Knowing the anatomical dimensions of the human organ and the size of the cell [4], we can estimate the number of cells *N*_*G*_ = *R*_0_*/k*_0_ in the cellular tissue of an adult organ.

In this work, we used the following estimates:

|  |  |
| --- | --- |
| Colon: | $N_G = 2.0 \cdot 10^8$ |
| Lung: | $N_G = 2.0 \cdot 10^{10}$ |
| Stomach: | $N_G = 2.4 \cdot 10^7$ |
| Breast: | $N_G = 5.0 \cdot 10^7$ |
| Prostate: | $N_G = 8.8 \cdot 10^6$ |

To compare the error values for the age distributions of different cancers for the obtained (optimized) parameters of the *Er* function, we calculate the relative approximation error as a percentage of the *D*_*max*_ (maximum incidence from the dataset):

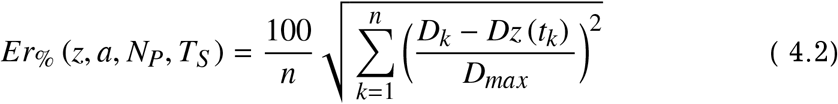

## 5. Results

The results are shown in Figures 5.1 and 5.2, where, for each type of cancer, the age distribution (open points in Figures 5.1 and 5.2) and theoretical age distribution curve (solid curves in Figures 5.1 and 5.2) are plotted.

**Figure 5.2.**
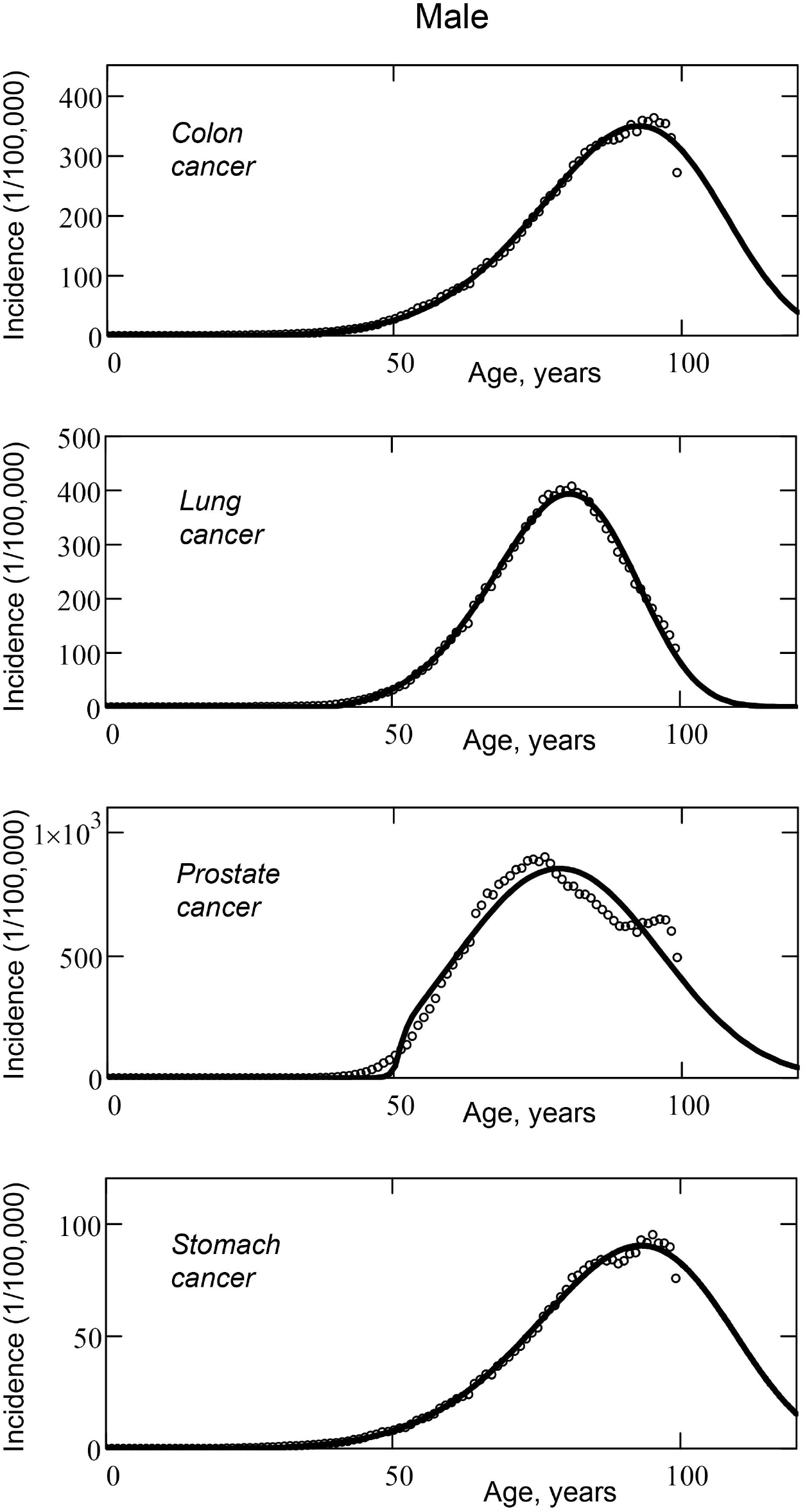
Age distribution of cancer incidence in men. The light points are real datasets; solid bold curves are approximation functions (mutational model of cancer).

The obtained parameters of the approximating function, at which the minimum approximation error is achieved, are shown in Table 5.1.

**Table 5.1.** Approximation parameters for the mutational model.

| **Female** | $z$ | $T_S$ | $a$ | $N_P$ | $Er$ | $Er\%$ |
| --- | --- | --- | --- | --- | --- | --- |
| Colon | 6 | 8.26 | $2.1 \cdot 10^{-1}$ | 11,291 | 0.46 | 0.16 |
| Lung | 5 | 24.11 | $2.8 \cdot 10^{-4}$ | 6,997 | 0.56 | 0.24 |
| Breast | 3 | 18.60 | $5.3 \cdot 10^{-6}$ | 22,146 | 1.25 | 0.31 |
| Stomach | 6 | 9.31 | $2.0 \cdot 10^{-1}$ | 1,746 | 0.13 | 0.29 |
| **Averaged** | **5.0** | **15.07** | **$1.0 \cdot 10^{-1}$** | **10,545** | **0.60** | **0.25** |

| **Male** | $z$ | $T_S$ | $a$ | $N_P$ | $Er$ | $Er\%$ |
| --- | --- | --- | --- | --- | --- | --- |
| Colon | 6 | 6.84 | $2.1 \cdot 10^{-1}$ | 13,818 | 0.65 | 0.19 |
| Lung | 5 | 25.68 | $2.5 \cdot 10^{-2}$ | 12,082 | 0.60 | 0.15 |
| Prostate | 3 | 35.62 | $7.2 \cdot 10^{-5}$ | 34,686 | 5.18 | 0.61 |
| Stomach | 6 | 2.91 | $1.3 \cdot 10^0$ | 3,754 | 0.17 | 0.19 |
| **Averaged** | **5.0** | **17.76** | **$3.8 \cdot 10^{-1}$** | **16,085** | **1.65** | **0.29** |

In the bottom row of Table 5.1, the parameter values averaged over four types of cancer (separately for women and men) are highlighted in bold.

The average number of mutations required to form a cancer is 5 mutations per cell for female and male cancer.

The average number of mutations per unit time is 1.0 · 10 ^−1^ mutations per year per cell for women and 3.8 · 10 ^−1^ mutations per year per cell for men.

The root mean square relative error of approximation by this mutation model is 0.25 percent for women and 0.29 percent for men (parameter *Er*_%_in table 5.1).

## 6. Discussion

Figures 5.1 and 5.2 clearly show that the age distributions of cancers in most cases (except for prostate cancer in men) are well approximated by the mutational cancer model. (light points of real data sets in men are closer to theoretical curves). The theoretical model best describes the age distribution of colon cancer in women and lung cancer in men – in the latter case, real data and theoretical curves practically coincide.

Average accuracy of approximation for four forms of cancer in women is slightly better than in men (bottom line, in bold, in Table 5.1, parameter *Er*_%_).

The mutational model used for data fitting allows us to determine the following parameters of cancer:

– the number of mutations per cell required for cancer to occur (*z* parameter from Table 5.1);
– the average number of mutations per unit time (year) per cell of the considered cell tissue (*a* parameter from Table 5.1);
– time lag in medical diagnosis of cancer (the time interval between the occurrence of cancer and its detection), (*T*_*S*_ parameter from Table 5.1);

Of the eight examined cancers (4 female and 4 male), only five cases show *T*_*S*_ (time of delay in diagnosis): lung cancer and breast cancer in women; lung cancer, prostate cancer and stomach cancer in men.

For other forms of cancer, with optimal parameters of the error function, the time *T*_*S*_ turns out to be negative, and this does not correspond to reality (in the case of negative time lag, we include in table 5.1 the last non-negative value of *T*_*S*_, which is obtained by sequential discrete increase in the number *z*). This fact may indicate the imperfection or inapplicability of the used mathematical model in this case.

In addition, the considered mutational model has other significant drawbacks.

First, it should be understood that the model describes the key events that happened to a biological cell, but the model says nothing about the causes of these events (the model does not state that these events are cellular mutations).

Second, the model does not tell us why the number of these events (or mutations) is five (bottom line of Table 5.1, in bold, *z* parameter) ? We do not know how we can practically measure this “magic” number in any other way than by looking at the age distribution of cancers. This indicates that we do not sufficiently understand the processes taking place in a biological cell.

Therefore, other mathematical models should also be considered to describe the age distribution of cancers. Perhaps we will try to present some of these models in one of the following articles.

Saint-Petersburg

2021

